# ProteinWeaver: A Webtool to Visualize Ontology-Annotated Protein Networks

**DOI:** 10.1101/2024.10.24.620032

**Authors:** Oliver Anderson, Altaf Barelvi, Aden O’Brien, Ainsley Norman, Iris Jan, Anna Ritz

**Author notes:** These authors contributed equally (Oliver Anderson, Altaf Barelvi).

## Abstract

Molecular interaction networks are a vital tool for studying biological systems. While many tools exist that visualize a protein or a pathway within a network, no tool provides the ability for a researcher to consider a protein’s position in a network in the context of a specific biological process or pathway. We developed ProteinWeaver, a web-based tool designed to visualize and analyze non-human protein interaction networks by integrating known biological functions. ProteinWeaver provides users with an intuitive interface to situate a user-specified protein in a user-provided biological context (as a Gene Ontology term) in five model organisms. Protein-Weaver also reports the presence of physical and regulatory network motifs within the queried subnetwork and statistics about the protein’s distance to the biological process or pathway within the network. These insights can help researchers generate testable hypotheses about the protein’s potential role in the process or pathway under study. Two cell biology case studies demonstrate ProteinWeaver’s potential to generate hypotheses from the queried subnetworks. ProteinWeaver is available at https://proteinweaver.reedcompbio.org/.

## 1 Introduction

Biological networks are essential tools for modeling and understanding complex biological systems [1, 2]. Protein-protein interaction (PPI) networks are biological networks that describe all the proteins and their molecular interactions within a biological system [3]. Species-specific PPI networks were first built from high-throughput protein interaction detection methods (such as yeast two-hybrid (Y2H) and affinity purification mass spectrometry (AP/MS) studies) [3] and are now generated from a diverse array of *in vitro, in vivo*, and *in silico* methods [4]. Graphs are powerful tools for modeling and analyzing the function and interactions of proteins within an organism [5, 6], and have been used to identify the differences between healthy and disease states in organisms [7]. PPI networks are valuable tools for analyzing molecular interactions across all domains of life, from viral to human tissue, and PPIs from non-human model organisms can be used to study human diseases [8].

In addition to PPI networks, gene regulatory networks (GRNs) are graphical representations of biological systems utilized for studying development, differences between healthy and diseased states, and other biological processes that are regulated by transcription factors (TFs) [9]. GRNs differ slightly from PPI networks in their graphical representation: there are directed interactions where TFs regulate a gene or gene product rather than undirected physical interactions. Although many existing tools represent PPI networks or GRNs graphically, many represent one type of network in isolation or fail to draw clear visual distinctions between the interaction types [10]. In addition, regulatory interactions and PPIs do not exist in isolation; physical and gene regulatory interactions coexist, forming intricate patterns or motifs. Physical interactions in these “mixed motifs” have been found to participate in regulatory interactions [11]. Methods to identify mixed motifs are lacking but can be useful in discovering clusters of proteins or gene products for further experimental analysis [10].

Various web tools have been developed to visualize molecular networks. STRING-DB is one of the most well-known network visualization tools, offering the ability to visualize subnetworks in different species by querying proteins or biological processes [12]. Other tools have been developed to visually explore specific pathways [13–15] or focus on individual species [16–19], and some require user-inputted experimental data or labeled genes [20–22]. See Supplementary Section S1 for more information about related web tools.

Despite the abundance of web servers for molecular network visualization, one may have a protein they are currently studying and want to know how it could participate in a particular pathway or function. To the best of our knowledge, no network visualization web server focuses on situating a protein of interest within a specific biological function. Tools requiring user input data make them difficult to use for general hypothesis generation. Additionally, some tools that run algorithms hide the details from the user, so it is unclear to the researcher how the subnetworks are selected for visualization. Thus, there is a need for a hypothesis generation tool for non-human model organisms that answers this simple question.

We present ProteinWeaver, a molecular interaction network visualization tool that generates subnetworks of physical and regulatory interactions based on a protein and a biological function of interest for non-human model organisms. Currently, ProteinWeaver supports a prokaryote (the Gram-positive bacterium *Bacillus subtilis subsp. subtilis str. 168*), a single-celled eukaryote (the brewer’s yeast *Saccharomyces cerevisiae S288C*), two morphologically-distinct invertebrates (the fruit fly *Drosophila melanogaster* and the nematode *Caenorhabditis elegans*), and a vertebrate (the zebrafish *Danio rerio*). ProteinWeaver characterizes biological functions with the Gene Ontology (GO), a collection of classifications for protein function (GO terms) [23, 24]. GO terms are a valuable tool for predicting protein function using PPI networks and can help situate a protein within a biological context [25]. ProteinWeaver generates a subnetwork that connects a protein of interest with proteins annotated to the specified biological context. The graphical interface is fast, visually intuitive, and does not require previous computational experience to use effectively.

ProteinWeaver offers two additional pieces of information to help situate a protein in the context of a biological process or pathway. It counts five different network motifs (one with PPI edges, another with regulatory edges, and three with a mix of PPI and regulatory edges) and provides enrichment scores for users to understand the expected motif count for an organism. ProteinWeaver also provides a quantitative measure that describes how close a protein is to proteins annotated to a biological process within the network. These additional features help provide context for the protein of interest and the surrounding molecular interactions for hypothesis generation.

## 2 Design and Implementation

### 2.1 Interaction Data

Currently, ProteinWeaver supports physical and regulatory interaction data for five non-human model organisms: *B. subtilis, C. elegans, D. melanogaster, D. rerio*, and *S. cerevisiae* (Table 1). For each organism, we collected experimental, text-mined, and database-validated protein and genetic interactions, as well as Gene Ontology (GO) annotations (Supplementary Section S2.1). All protein-protein and regulatory interactions are linked to evidence sources, such as PubMed or STRING-DB references [12].

**Table 1:** Organisms and their associated interaction networks hosted by ProteinWeaver. Nodes represent proteins and their encoding genes. PPI: protein-protein interaction. GRN: gene regulatory network.

| Organism | Nodes | PPI Edges | GRN Edges | GO Terms | GO Annotations |
| --- | --- | --- | --- | --- | --- |
| *B. subtilis* | 3,163 | 6,441 | 5,634 | 3,681 | 78,015 |
| *C. elegans* | 4,106 | 13,915 | 78,223 | 7,868 | 202,845 |
| *D. melanogaster* | 12,823 | 233,054 | 17,530 | 11,774 | 492,331 |
| *D. rerio* | 16,606 | 45,003 | 25,960 | 8,321 | 133,619 |
| *S. cerevisiae* | 7,644 | 164,432 | 237,315 | 8,300 | 328,186 |

The relationships among proteins and GO terms are represented as a connected graph with two types of nodes (proteins and GO terms) and three types of edges that capture undirected physical interactions among proteins, directed regulatory interactions among proteins, and directed GO term annotations between GO terms and proteins (Figure 1 and Supplementary Section S2.2). We add the directly annotated protein-GO Term pairs into the graph and infer annotations between proteins and more general GO terms (purple edges in Figure 1). ProteinWeaver uses the Neo4j Graph Database to store and query the graph. See Supplementary Section S2.3 for more information about Neo4j and Supplementary Section S2.5 for information about the ProteinWeaver development stack.

**Figure 1:**
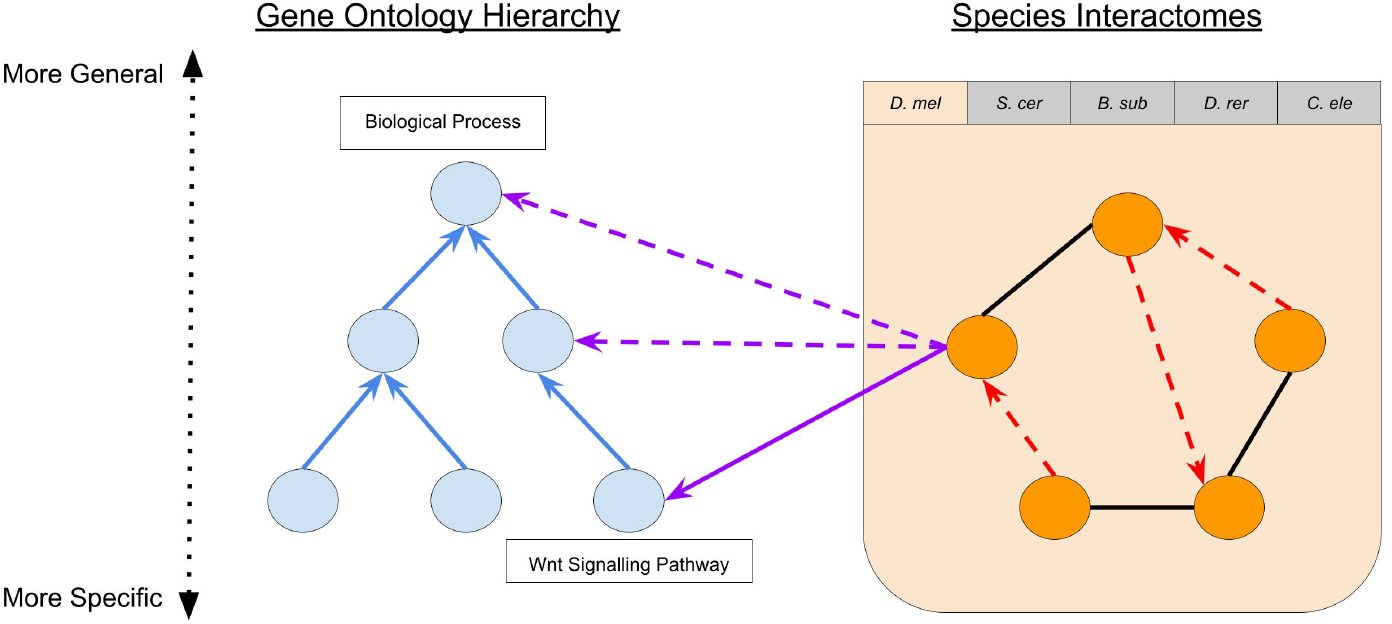
Graph representation of Proteins and Gene Ontology terms. Organism-specific networks contain protein-protein interaction edges (black solid lines), regulatory edges (red dashed arrows), and proteins (orange circles). The GO term hierarchy is represented via the blue nodes and directed blue solid edges. Proteins directly annotated to GO terms are indicated by purple solid arrows, and purple dashed arrows indicate inferred annotations.

### 2.2 Queried Subnetworks

The fundamental feature of ProteinWeaver is the ability for a user to enter a query protein *s* and a GO term *t* and visualize the connections from *s* to proteins annotated to *t* (Figure 3A). Users specify the size of the subnetwork returned by selecting a small integer *k* (which typically ranges between 5 and 25). ProteinWeaver generates these subnetworks in two different ways: in “K Unique Paths” we calculate the *k*-shortest paths from *s* to nodes annotated to *t* using Yen’s *k*-shortest paths algorithm [26], and in “K Unique Nodes” we calculate the *k* nodes annotated to *t* that are the closest to *s* using a breadth-first search from *s* to reachable nodes annotated to *t*. For both algorithms, increasing the value of *k* increases the size of the visualized network. More information about the algorithms used for subnetwork generation can be found in the supplementary Section S2.4. When we visualize the subnetwork, we do not include the GO term *t*, thus ending the path at the proteins annotated to *t* (e.g., the blue nodes in Figure 2).

**Figure 2:**
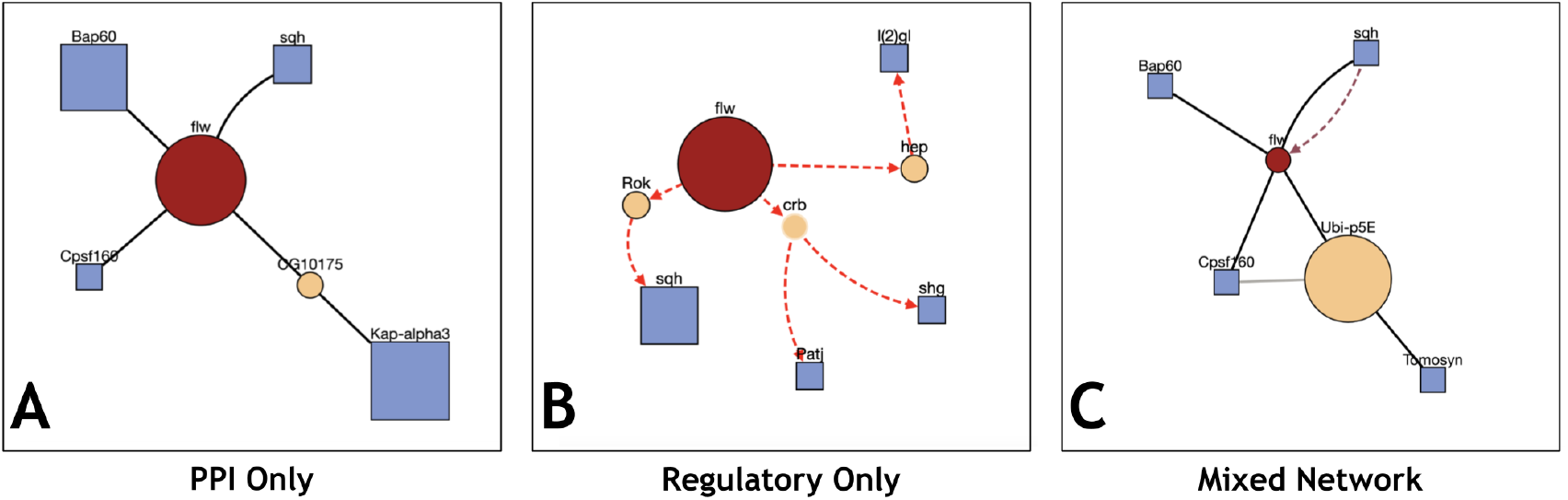
ProteinWeaver results from a “K Unique Nodes” query connecting flapwing (flw) to *k*=4 nodes annotated to”myosin binding” (GO:0017022) *D. melanogaster* with (A) physical interactions, (B) regulatory interactions, and (C) both physical and regulatory interactions. Dashed directed edges indicate regulatory relationships, solid edges indicate undirected physical relationships; see Fig 3B for a full legend describing node and edge types.

**Figure 3:**
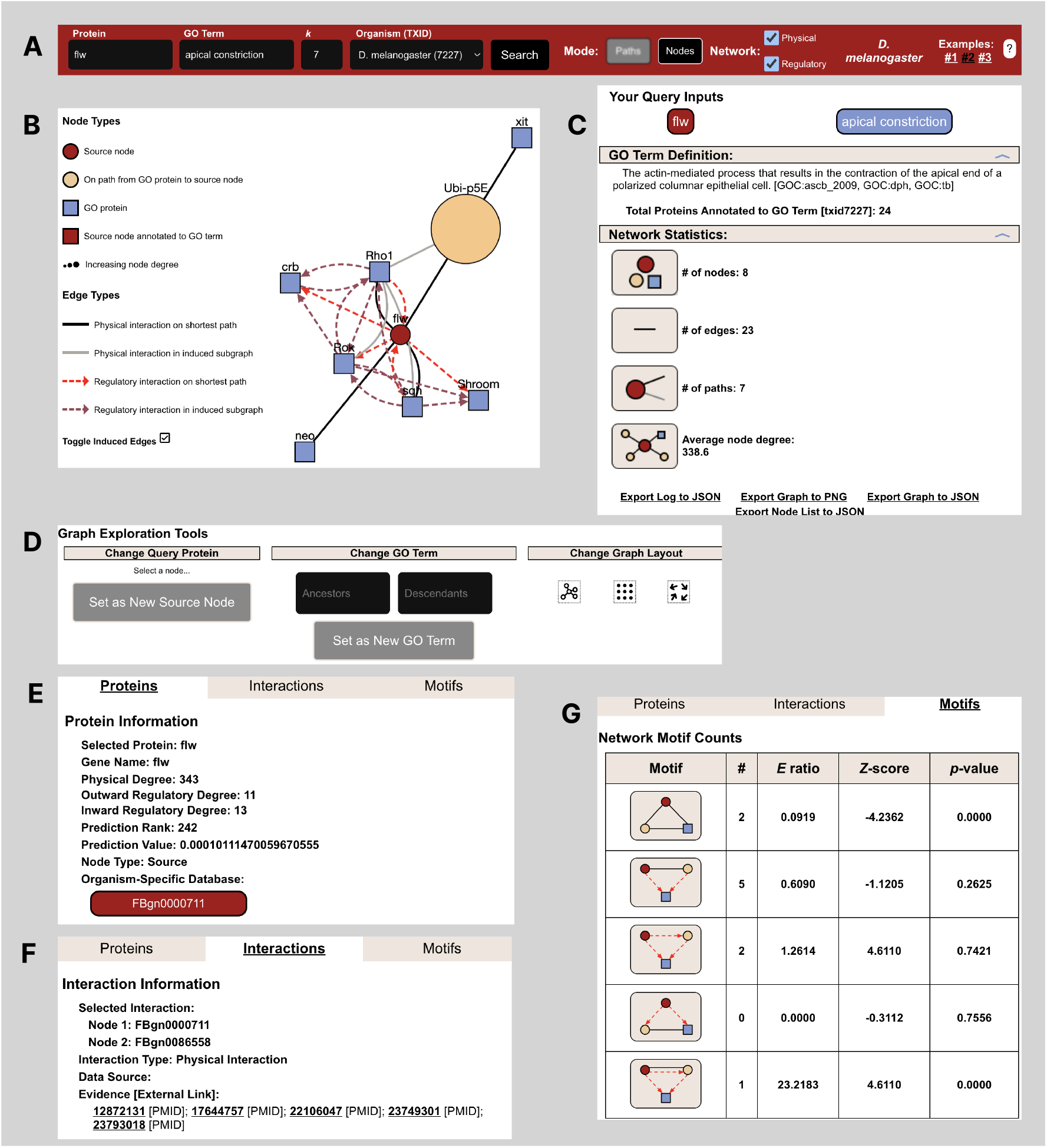
The ProteinWeaver interface. A) Users enter their protein and GO term of interest, a small integer *k*, and the species in the top panel and click “Search”. B) The subnetwork is shown via `react-cytoscapejs`’s plugin. The legend section details the different node and edge types. C) The sidebar includes links to protein and GO term data, basic network statistics, and the ability to export the session. D) The graph exploration section lets users rearrange the network layout and update the query by selecting a new query node or traversing the gene ontology. E-G) The tabbed window shows information about the user-selected protein (including the protein’s rank relative to the GO term), information about the user-selected interaction, and the motifs found in the subnetwork.

ProteinWeaver can return subnetworks consisting of physical, regulatory, or mixed physical and regulatory interactions. The same query run with different interaction types produces different subnetworks (Figure 2). All three subnetworks included common GO-annotated proteins, such as sqh, but each also identified unique GO-annotated proteins, offering different contexts for the source protein based on interaction type. Researchers can thus choose to query specific networks of interest or combine different interaction types and query modes to explore their protein of interest in a broader biological context.

### 2.3 ProteinWeaver Interface

ProteinWeaver allows users to explore networks for non-human model organisms by inputting a protein, GO term, and a size parameter (*k*), specifying whether to traverse a protein-protein interaction (PPI) network, gene regulatory network (GRN), or a mix of both (Figure 3A). The induced subnetwork gives users a detailed view of how nodes are interconnected within the organism (Figure 3B). Users can navigate the network by selecting GO terms from the hierarchy or changing the queried protein to any protein within the subnetwork, fostering hypothesis generation based on the species’ interactome (Figure 3D).

For each query, ProteinWeaver provides comprehensive details, including annotation links for the queried protein and GO term, GO term definitions, and links to organism-specific databases (Figure 3C & Figure 3E). The tool also includes a statistics section showing graph statistics, GO term annotation confidence scores, and mixed motif data (Figure 3C, Figure 3E, & Figure 3G). To support reproducibility, queries can be saved as hyperlinks or exported as PNGs, JSON files, or Cytoscape objects for further analysis.

## 3 Results

### 3.1 Mixed Motif Enrichment

Many biological processes involve both regulatory and physical interactions. Consequently, representing these networks separately can obscure the complete functional context of a protein of interest [10]. Physical interactions involving transcription factors (TFs) have also been shown to effectively predict long-range enhancer-promoter interactions [11]. Therefore, a method for identifying mixed regulatory-physical interaction clusters, or “mixed motifs,” is valuable for researchers seeking a comprehensive understanding of the regulation of specific proteins or processes.

ProteinWeaver displays statistics for five network motifs that were found to be significantly over-represented in a *S. cerevisiae* mixed PPI-Regulatory network [10]. These 3-node motifs consist of three mixed motifs and two network-specific motifs enriched in the yeast interactome and carry explainable biological significance. The PPI-specific motif, “protein clique” (Figure 4A), often represents three proteins working together in a multi-protein structural or functional unit. The other network-specific motif, “feed-forward loop” (Figure 4B), consists of two TFs, one of which regulates the other, regulating a third protein or gene and is a canonical regulatory motif [27] found to be enriched in *S. cerevisiae, E. coli*, and *C. elegans* transcriptional networks [28–30].

**Figure 4:**
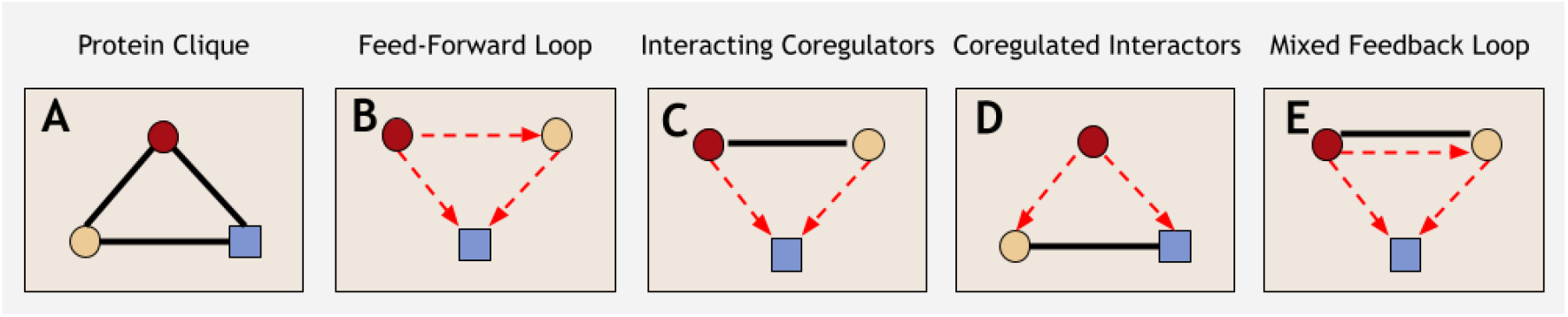
Mixed network motifs identified by ProteinWeaver; adapted from Yeger-Lotem et al. [10]. Black lines indicate physical interactions. Red arrows indicate regulatory interactions. Motifs can contain multiple protein types.

In addition to recognizing well-known PPI and gene regulatory motifs, ProteinWeaver can identify mixed network motifs. The first, “interacting coregulators” (Figure 4C), represents two proteins that physically interact and regulate the same gene. The second mixed motif, “coregulated interactors” (Figure 4D), represents two physically interacting proteins that the same TF regulates. The final mixed motif, the “mixed feedback loop” (Figure 4E), consists of a feed-forward loop where the two TFs physically interact. Physical interactions between TFs can be regulatory active themselves [11]. Thus, identifying these motifs is valuable for researchers exploring the regulatory implications of physical binding events. Additionally, proteins that work together are often coregulated. Therefore, proteins identified in a coregulated interactor motif may be more likely to have functional impacts on each other [10]. It is important to note that only by using a mixed network can ProteinWeaver capture these last three motifs accurately, as a purely regulatory network would incorrectly classify a mixed feedback loop as a feed-forward loop.

The background distribution of motifs in the *B. subtilis, D. melanogaster*, and *D. rerio* graphs are relatively similar, with all of them having a large number of Protein Cliques relative to the other four motifs (Figure 5). *C. elegans* has a much larger relative proportion of Feed-Forward Loops than the other species and has more evenly distributed motifs than *B. subtilis, D. melanogaster*, and *D. rerio. S. cerevisiae* has the most uniformly distributed network, with many instances of all five motifs being found at a relatively high rate. *S. cerevisiae* being the most uniformly distributed may reflect a bias since the motifs identified here are based on motifs found enriched in a yeast network [10]. For a queried subnetwork, ProteinWeaver counts the number of each type of motif in the network (Figure 3G). Enrichment scores, *Z*-scores, and *p*-values are calculated compared to the species-specific background graph; see Supplementary Section S3.1 for more information.

**Figure 5:**
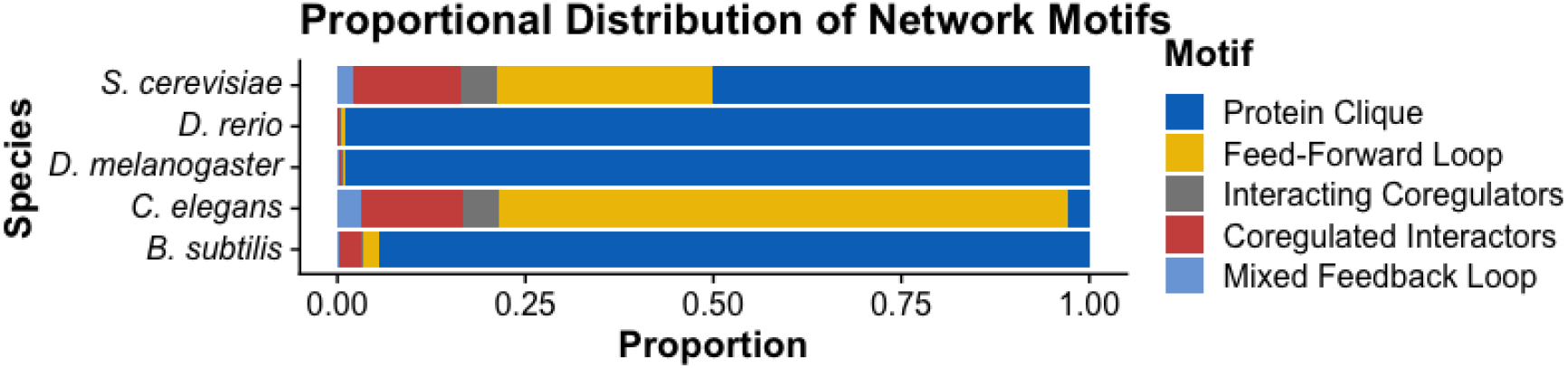
Proportion of motifs by species in ProteinWeaver. For a detailed breakdown of the motifs by species, see Supplementary Section S3.2

### 3.2 GO Term Annotation prediction

In a typical ProteinWeaver query, the query protein *s* is not associated with the queried GO term *t*, and the goal is to connect the query *s* to proteins annotated to *t*. Sometimes, the queried protein *should* be considered part of the GO term, but there is not yet evidence in the Gene Ontology to properly annotate the protein. To provide the user additional context about whether the query is likely to be associated with the GO term of interest, we used a random walk approach to assign a confidence score about whether a query protein *s* is “near” GO-term annotated proteins. The random walk scoring function runs personalized PageRank [31], restarting from proteins annotated to the GO term *t* with a damping factor *α* = 0.7.

We rank the query node *s* according to the final visitation probabilities of the random walk; see Section S4 for more details.

We first compare the random walk approach (which we call **RandomWalk**) to three other prediction scores based on neighbor overlaps in the graph:

#### Degree

Rank *s* by its degree in the original graph *G* (larger is better). This approach does not account for the GO term *t*, and it is used to assess degree bias in our evaluation.

#### One-Hop GO Overlap

Rank *s* by the number of *s*’s neighbors that are annotated to GO term *t* (larger is better).

#### Hypergeometric Distribution

Rank *s* by the hypergeometric distribution *p*-value of observing the number of *s*’s neighbors annotated to GO term *t*, adjusted by the size of the GO term *t* and the degree of *s* (smaller is better).

See Supplementary Section S4.1 for more details about these comparator methods.

We assessed the four methods by generating a dataset of 1,000 positives, which were selected by randomly choosing a protein-GO term edge and modifying *G* to remove that specific edge. We expect these nodes to have high confidence in their membership with the GO term. For every positive (protein-GO Term pair), we selected 100 negative proteins by identifying proteins that are (1) near the positive protein when considering PPI and regulatory edges only, (2) are not connected to the positive GO term, and (2) have approximately the same degree as the positive node. See Section S4.2 for more information about positive and negative sampling and the evaluation pipeline. We generated these datasets for each species and plotted precision-recall and ROC curves (Figures 6 and S7).

**Figure 6:**
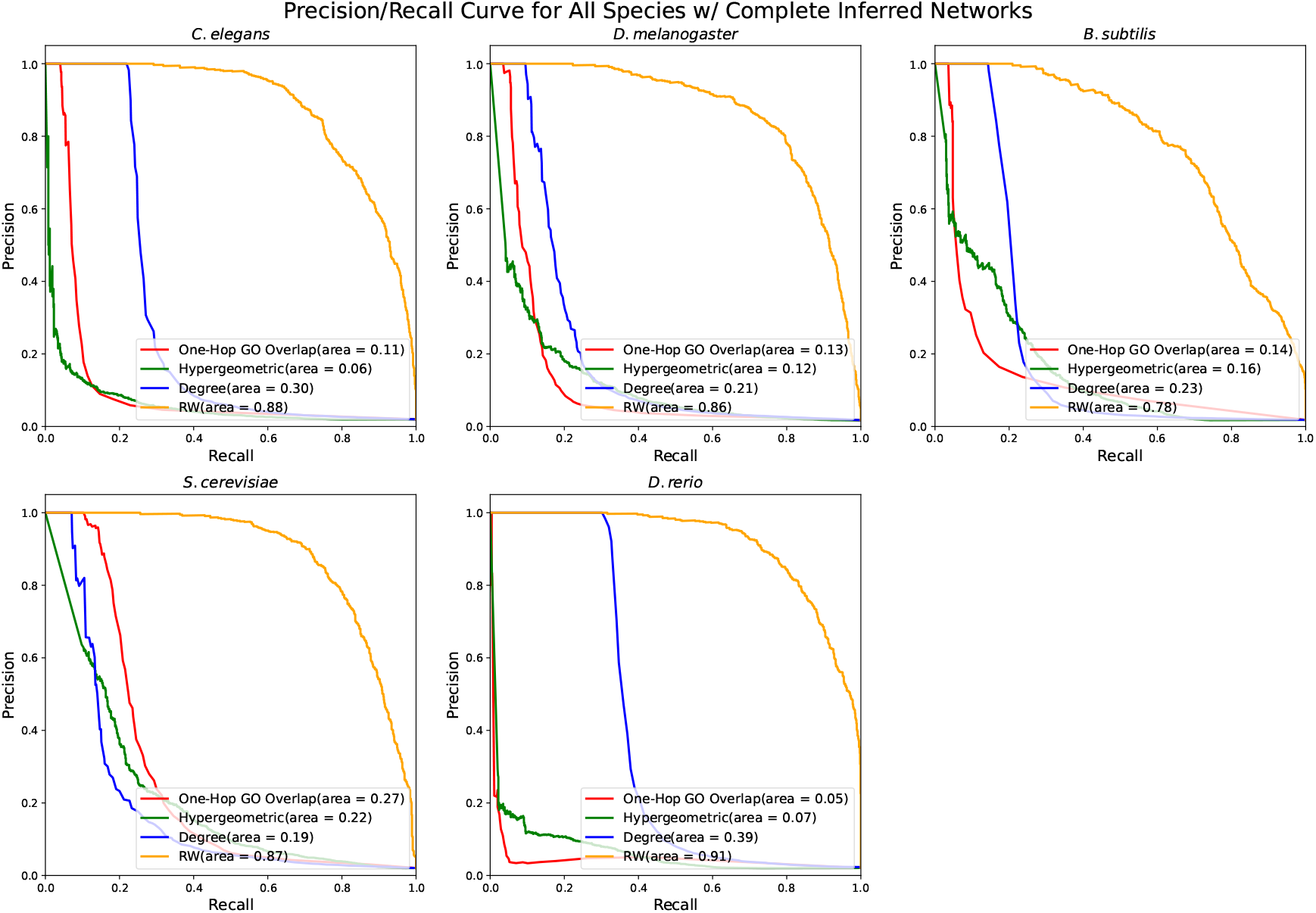
Precision-Recall curves of four annotation prediction methods across all species with inferred networks. Examples were sampled at a 1:100 positive-to-negative ratio.

We found that RandomWalk had a nearly perfect ROC AUC across all species (Figure 6. RandomWalk also had the highest precision at varying recall values and it drops off precision at higher recall values. The rest of the methods performed considerably worse than RandomWalk in the ROC AUC analysis. Between the methods, their ranks varied in different species analysis for both the inferred and non-inferred networks. For example, the degree method was ranked second in the ROC value for *C. elegans*, however was ranked 4th for the yeast dataset. The Precision/Recall values for the One-Hop GO Overlap and Degree methods noticed an increase when using the inferred networks. The significance of this increase varied among the methods and which species. For example, the degree method noticed the largest Precision/Recall increase in the *C. elegans* dataset. The Hypergeometric method, however, did not show any noticeable difference between the Precision/Recall values across all the species when using the inferred and non-inferred networks. This could be because adding inferred protein-GO edges did not affect the overall hypergeometric equation used to calculate their scores. We also ran these methods on the graphs with only directly-annotated networks; while the relative ordering of the three comparator methods changed, the random walk approach remained superior (Supplementary Section S4.3). Furthermore, we tested the performance of RandomWalk on four different background networks (by selecting different subsets of edge types) and found it had little effect on the ROC and PR curves (Supplementary Section S4.4).

### 3.3 Case Studies

To illustrate the potential of ProteinWeaver in generating hypotheses, we describe two case studies from the recent literature.

#### D. melanogaster

The protein Eb1 is part of a group of the end-binding proteins family responsible for microtubule plus end growth [32]. Microtubules are important polymers in all eukaryotes that play the role of changing a cell’s shape, division, and transport [33]. Microtubules interact with microtubule-associated proteins (MAPs) to regulate microtubules in the cell [33]. In a 2021 paper, the gene Eb1, along with Tau and XMAP215/Msps were discovered to cooperate independently in the axon of Drosophila to regulate microtubule polymerization and bundle formation [34]. Furthermore, in a 2013 paper, Eb1 was noted to be important in apicobasal microtubule bundle formation and epithelial elongation [35]. However, in the Gene Ontology database, Eb1 has not yet been annotated to microtubule bundle formation in Drosophila. We queried the connections between Eb1 and microtubule bundle formation to generate a visualization of Eb1’s connection to other microtubule bundle-related proteins (Figure 7A). From the visualization, Eb1 is connected to important microtubule-associated proteins such as Short stop (Shot), which binds actin and microtubules, and Ncd, which produces a minus-end-directed kinesin microtubule motor protein [36, 37]. Additionally, ProteinWeaver identifies the protein Smt3, which is linked to microtubule-associated proteins but lacks direct annotation for microtubule bundle formation. Smt3 is part of the Small Ubiquitin-like Modifier (SUMO) family, which plays a key role in the post-translational modification of target proteins through SUMOylation. Notably, proteins involved in mitotic spindle formation and organization have been reported as substrates for SUMOylation [38]. Among the 12,800 fly proteins not annotated to microtubule bundle formation, Eb1 was ranked 40th according to the RandomWalk prediction.

**Figure 7:**
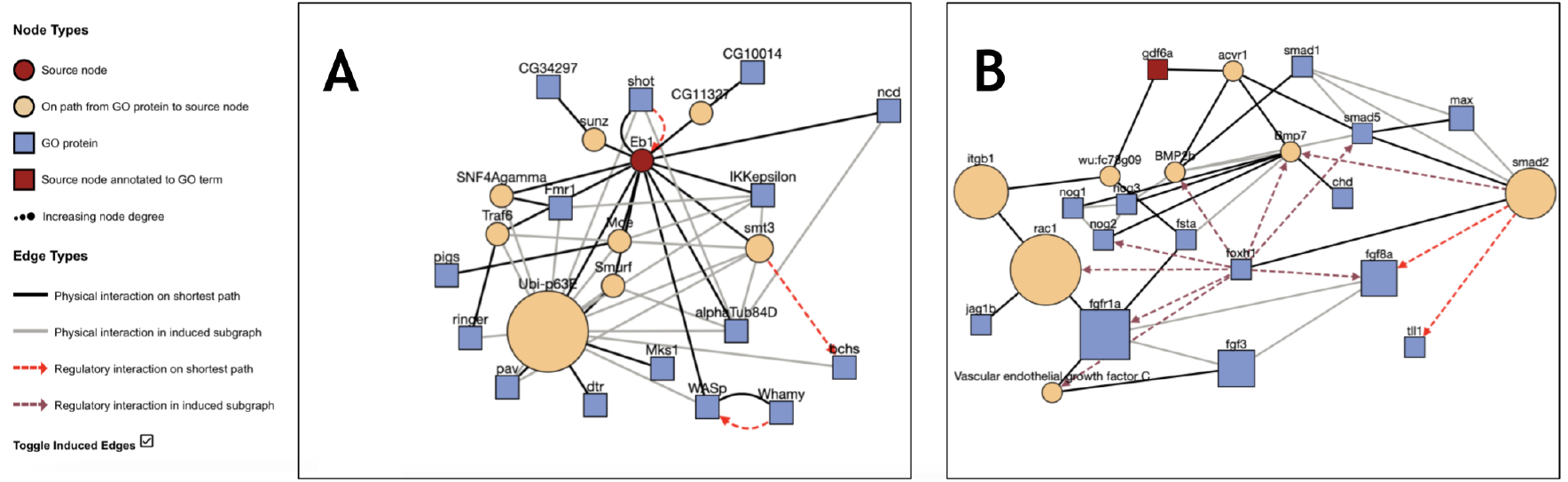
ProteinWeaver showcasing subnetworks from A) Eb1 to “microtubule bundle formation” (GO:0001578) in *D. melanogaster* and B) Gdf6a to “dorsal/ventral pattern formation” (GO:0009953) in *D. rerio*. Both queries can be found through these URLS respectively, https://tinyurl.com/proteinweaver-study-1 and https://tinyurl.com/proteinweaver-study-2

#### D. rerio

Bone morphogenic proteins (BMPs) have been shown to function in a variety of processes in animals, including patterning and differentiation of tissues, establishing cell polarity, maintaining organ homeostasis, and responding to injuries [39]. The BMP pathway in zebrafish is regulated in part by the Smad family of transcription factors [40]. Further, BMPs have been found to regulate Smads through phosphorylation via a receptor complex [39]. Previous work has shown that knocking out a specific BMP, *gdf6a*, blocks retinal Smad phosphorylation, indicating that Gdf6a is involved in regulating Smad proteins in the retina [41]. We wanted to visualize the connection between the BMP family member, Gdf6a, and other proteins linked to dorsal/ventral patterning to investigate how Smad transcription factors might integrate BMP with other signaling pathways to promote D/V pattern formation.

When querying Gdf6a with the GO term “dorsal/ventral pattern formation,” ProteinWeaver identified a receptor protein serine/threonine kinase ACVR1 that connects Gdf6a to the proteins annotated to dorsal/ventral pattern formation (Figure 7B). ACVR1 interacts with Gdf6a, Bmp7a, Bmp2B, and Smad5 and is involved in left/right patterning in mice development [42]. These proteins interact with more Smad family transcription factors; Smad2 is particularly interesting because it is not annotated to the queried GO term “dorsal/ventral pattern formation,” and Smad 1/5/9 are usually associated with this patterning in zebrafish [43]. However, Smad2 has been linked to dorsal mesoderm specification in *Xenopus*, and mutants have shown that Smad2 is essential in early embryo patterning events in mice [44]. Among the almost 16,000 nodes not annotated to “dorsal/ventral pattern formation,” ProteinWeaver ranks Smad2 as the 12th most likely to be annotated to the GO term.

## 4 Availability and Future Directions

ProteinWeaver is licensed under GNU General Public License v3.0 and is available at https://proteinweaver.reedcompbio.org/. All data, resources, scripts, and website source code are available at https://github.com/Reed-CompBio/protein-weaver/. Protein Prediction data is available at https://github.com/Reed-CompBio/protein-function-prediction.

ProteinWeaver is designed to be an efficient, user-friendly, and reproducible tool for GRN and PPI network analysis. ProteinWeaver provides researchers with novel network querying capabilities of protein interactions, regulatory interactions, or a combination of interaction types for five species. Mixed motif identification and GO term annotation predictions serve as additional resources for hypothesis generation and network exploration in these non-human model organisms.

We note that our approach to predict GO term annotations is a different problem than that of determining protein function, which is a longstanding challenge that has seen explosive improvements with structural alignments provided by deep learning models such as AlphaFold [45]. Here, we aim to use the physical, regulatory, and GO-annotated relationships to assess how “near” a protein is to a GO term without additional information such as protein sequences, domains, or structure.

Currently, ProteinWeaver supports five non-human model organisms but plans to expand to include more. ProteinWeaver wants to incorporate more prokaryotic model organisms such as *E. coli* and eukaryotic plant species like *Arabidopsis thaliana*. Current limitations are related to the lack of freely available physical and regulatory network data for specific species. We are also working to provide more context about the local subnetwork structure in relation to the GO-annotated proteins and other nearby interacting molecules beyond the current basic statistic. This context will come in the form of expanded network statistics, such as centralities and clustering coefficients, modified to account for proteins annotated to the GO term of interest.

In addition, ProteinWeaver aims to improve its querying capabilities by allowing users to search with multiple proteins of interest or multiple GO terms of interest. This facilitates network exploration for researchers interested in how several proteins interact within a specific biological process or how multiple biological processes may connect.

ProteinWeaver aims to maintain an intuitive and straightforward user interface while providing dense information. Striking the right balance between these goals poses a challenge, and further enhancements are made continually to optimize user experience while maintaining informativeness. ProteinWeaver invites collaboration with the scientific community for feature development. Open discussions and feedback sessions will guide the implementation of features aligned with user needs and advancements in biological research.

In conclusion, ProteinWeaver facilitates the understanding of molecular interactions and their roles in biological contexts, even for those without computational expertise. Its intuitive interface and GO term integration address challenges researchers face in situating proteins within a specific biological context. ProteinWeaver is positioned to be a useful tool for hypothesis generation and biological interaction exploration for researchers studying non-human model organisms.

## 5 Acknowledgements

We thank Larry Zeng for his help with the web server and the tech stack architecture. We also thank Derek Applewhite, Kara Cerveny, and Shivani Ahuja for their collaborative discussions. This work was funded by NSF-DBI-1750981 (awarded to AR).

