## Supplementary Information for "ProteinWeaver: A Webtool to Visualize Ontology-Annotated Protein Networks"

#### S1 Existing Tools for Visualizing Molecular Networks

STRING-DB is one of the most well-known network visualization tools, offering the ability to visualize subnetworks in different species by querying proteins or biological processes [1]. Another commonly used PPI network visualization tool, GeneMANIA [2], uses additional functional genomic data to generate subnetworks. Some visualization tools are focused on specific subsets of the larger networks, such as signaling pathway visualization tools like SignaLink [3] and KEGG [4]. Many visualization tools are tailored for individual species, such as BioPlex [5] and GenePlexus [6] for humans, the Drosophila Interactions Database [7] for flies, and SubtiWiki for *B. subtilis* [8].

Finally, other network visualization tools such as KeyPathwayMiner [9] and NetworkAnalyst [10] require user data as input and/or run specialized algorithms. Although the network visualization capabilities of these tools are comprehensive, none of these existing tools allow users to search by a protein and specific pathway and get a subnetwork grouped by both their function and interactions with the protein of interest.

### S2 ProteinWeaver

#### S2.1 Data Sources

ProteinWeaver contains physical and regulatory information for five different species.

***B. subtilis***: For the Gram-positive bacterium *B. subtilis*, we exported the "Interaction" dataset from SubtiWiki (v4) [8] and merged it with the experimental, text-mined, and database-validated interactions from the "Physical Links" dataset from STRING-DB (v12) [1] to generate the PPI network. To generate the GRN, we exported the "Regulatory" dataset from SubtiWiki (v4). In addition, GO annotations were obtained from the "Gene" dataset from the SubtiWiki (v4) exports page and the QuickGO annotation service from the EBI [11]. UniProt IDs [12] were assigned to each of the proteins using the AmiGO 2 search function on the Gene Ontology web server [13].

***C. elegans***: For the nematode *C. elegans*, we obtained the PPI network through WormBase (v18), and filtered to only "physical interactions" [14]. To filter further, we only included interactions where both proteins were UniProt verified [12]. The regulatory interactions were exported from TFLink's database [15]. The GO term association data were also obtained through the WormBase database, and the UniProt namespace mapping service was used to convert between WormBase IDs and UniProt IDs [12].

***D. melanogaster***: For the fruit fly, we exported an existing fly PPI network from a compilation of six fly interaction databases [16]. The majority of the interactions in the PPI network come from the Drosophila Interactions Database (DroID) [7]. To generate the GRN, we exported the "Genetic Interaction Table" dataset from FlyBase (v2024_03) [17, 18]. To annotate the GO terms to proteins for *D. melanogaster*, we exported the Gene Association file from FlyBase and merged it with a dataset generated from the EBI's QuickGO annotation service [11]. UniProt IDs were assigned from FlyBase IDs to each protein using the UniProt name mapping service [12].

***D. rerio***: We downloaded protein aliases and interaction data from STRING-DB (v12) and filtered the data to experimental, text-mined, and database-validated validated interactions [1]. In addition, we scraped data from the PSICQUIC database [19] using a Python script. We pre-processed the XML into JSON using a JavaScript script and converted the JSON into a CSV format using an online converter [20]. In addition, we exported UniProt identifiers for *D. rerio* from the UniProt ID mapping service [12]. During this process, we found many of the interactor identifiers scraped from PSICQUIC to be obsolete. As a result, we discarded interactions where the proteins were not UniProt verified. For the GRN, we exported the regulatory interactions from TFLink [15]. Finally, we employed the EBI's QuickGO service to gather the GO annotations for *D. rerio* [21].

***S. cerevisiae***: For brewer's yeast, we chose the taxon ID 559292 since it appeared to be the strain with the most data available. We sourced our PPI network from BioGRID [22] while the regulatory interactions were exported from TFLink's database [15]. For GO term annotation data, we used QuickGO to find all reviewed annotations [21]. The UniProt namespace mapping service was used to convert between BioGRID IDs and UniProt IDs [12].

| Organism | Total | GRN-Only | PPI-Only | Shared |
|---|---|---|---|---|
| *B. subtilis* | 3,163 | 1,230 | 484 | 1,449 |
| *C. elegans* | 4,106 | 1,083 | 411 | 2,612 |
| *D. melanogaster* | 12,823 | 1,322 | 7,905 | 3,596 |
| *D. rerio* | 16,606 | 10,168 | 2,833 | 3,605 |
| *S. cerevisiae* | 7,644 | 858 | 1,092 | 5,694 |

Table S1: Breakdown of nodes for each organism in ProteinWeaver. GRN-Only nodes are only found in the regulatory network and do not have any interactions in the PPI network. PPI-Only nodes do not have any interactions in the GRN. Shared nodes have both genetic and regulatory interactions that they are associated with.

In addition to importing the interactions between proteins and the GO terms annotated to the proteins, we also imported the current Gene Ontology from the Gene Ontology web server (2024-07-17) [23] to facilitate the exploration of protein networks through the GO hierarchy. All the raw data and code is available on GitHub (see Data Availability).

| Organism | Total | Direct | Inferred |
|---|---|---|---|
| *B. subtilis* | 78,015 | 14,384 | 63,631 |
| *C. elegans* | 202,845 | 42,898 | 159,947 |
| *D. melanogaster* | 492,331 | 98,799 | 393,532 |
| *D. rerio* | 133,619 | 29,065 | 104,554 |
| *S. cerevisiae* | 328,186 | 69,760 | 258,426 |

Table S2: GO term annotations for each organism in ProteinWeaver. Annotations to ancestral GO terms were inferred from the direct annotations using the GO term hierarchy.

The scripts used to process and generate these datasets are available at the ProteinWeaver GitHub repository https://github.com/Reed-CompBio/protein-weaver/.

#### S2.2 Underlying Graph Representation

ProteinWeaver structures the interaction and GO term data into one unified graph $G = (V, E)$ with several node types and relationships (Figure 1 in the main text). The node set $V$ consists of all proteins from all species and the set of GO terms with at least one annotation for one of the proteins. There are four types of edges in the edge set $E$ that connect these nodes:

1. Undirected protein-protein interactions (*ProPro edges*).
2. Directed gene-regulatory interactions among protein products (*Reg edges*).
3. Directed protein-GO term edges that indicate GO annotations (*ProGo edges*).
4. Directed edges between GO terms that indicate the ontology hierarchy (*GoGo edges*).

Additionally, we infer indirect protein-GO term annotations from directly annotated descendants, which is common practice due to the transitivity principle [23, 24]. Since GO terms are hierarchical, we infer *ProGo* edges from specific GO terms to more general terms to capture the complete range of annotations for each protein [23, 24]. These relationships form a single (weakly) connected component $G$ (Figure 1 in the main text).

#### S2.3 Graph Database

ProteinWeaver leverages the Neo4j Graph Database with the Cypher query language for data management and querying (Figure S1). As a graph database, Neo4j can handle many-to-many relationships and large annotated networks more efficiently than traditional relational databases and is easier to use than other graph databases [25, 26]. Neo4j's native graph database structure is built for large and complex networks, and the database has several built-in graph algorithms accessible with Cypher.

The protein nodes contain properties with the name, unique database identifier, and species of origin. The GO term nodes contain properties with their name, ID, definition, and GO aspect. The *ProPro* and *Reg* interactions are recorded among proteins within each species. The *ProGo* relationships represent functional annotations linking proteins to GO terms. Proteins are annotated to the most specific GO term available. The *ProGo* edges have a property describing the direct annotation or an "inferred from descendant" tag, distinguishing between the most specific annotations and inferred ones. The *GoGo* edges have a property that describes the ontology between the two terms.

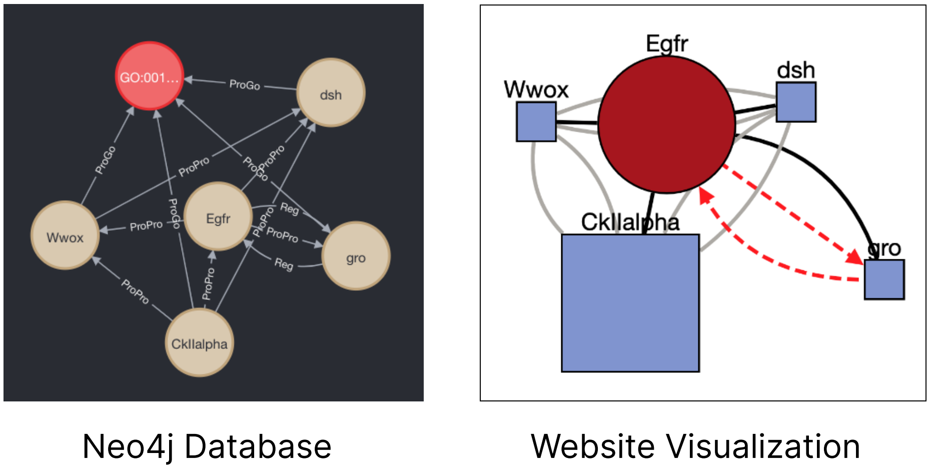

Figure S1: ProteinWeaver performing a "K Unique Paths" query. Neo4j represents protein nodes in beige and GO term nodes in red. Paths are generated connecting the protein of interest, Egfr, and the GO term "Wnt signaling pathway" (GO:0016055) with $k = 4$ in *D. melanogaster*. ProteinWeaver parses the query and only displays the protein nodes as a subnetwork to users.

#### S2.4 Path-based Algorithms

**K Unique Paths:** In "K Unique Paths" mode, ProteinWeaver generates subnetworks with a user-specified number of paths to proteins annotated to the GO term of interest. ProteinWeaver takes a user-defined protein, GO term, and $k$ and feeds the input into Neo4j's Yen's K-Shortest Paths algorithm to generate the graphical response [27]. The Yen's K-Shortest Path algorithm works similarly to Djikstra's Shortest Path with an additional $k$ parameter [28]. The $k$ parameter is the user-defined number of paths they want to display. ProteinWeaver finds the $k$-shortest paths from the protein of interest to proteins annotated to the GO term of interest. One limitation of this method is that it does not mark previously traversed edges, resulting in highly condensed graphs as

$k$ increases. To allow users to manually add more proteins annotated to their GO term of interest and create more diverse subnetworks, we developed the "K Unique Nodes" mode.

**K Unique Nodes:** We also offer a "K Unique Nodes" mode, which allows users to generate subnetworks that connect the query node $s$ to the closest $k$-nodes annotated to the GO term $t$. ProteinWeaver uses Neo4j's All Shortest Paths algorithm to efficiently compute all possible shortest paths from $s$ to reachable nodes in the graph [28]. We then order the nodes annotated to GO term $t$ based on the shortest path length from $s$ and display the paths that reach the first $k$ GO-annotated nodes.

##### S2.5 Tech Stack

The ProteinWeaver user interface is built with React JavaScript (Figure S2). React provides a scalable, user-friendly interface for responsive and intuitive web applications. We built the backend infrastructure using Node.js and Express, a web application framework for Node.js. This backend allows fast and lightweight handling of server-side logic and API calls. To visualize the PPI networks, ProteinWeaver employs the plugin `react-cytoscapejs` within the React framework. The plugin integrates the React framework with the Cytoscape.js biological network visualization tool, acting as a bridge from the user input to the Cytoscape user interface. ProteinWeaver also uses an additional Python backend server via Flask, which is mainly used to calculate GO term annotation prediction statistics. ProteinWeaver is hosted on the DigitalOcean platform,a secure and efficient environment for deploying web applications that provides a fast, stable web server. The codebase is also actively maintained and version-controlled on GitHub to facilitate open access and collaboration with others (see Data Availability). All software used to build ProteinWeaver is acknowledged on the website's About page.

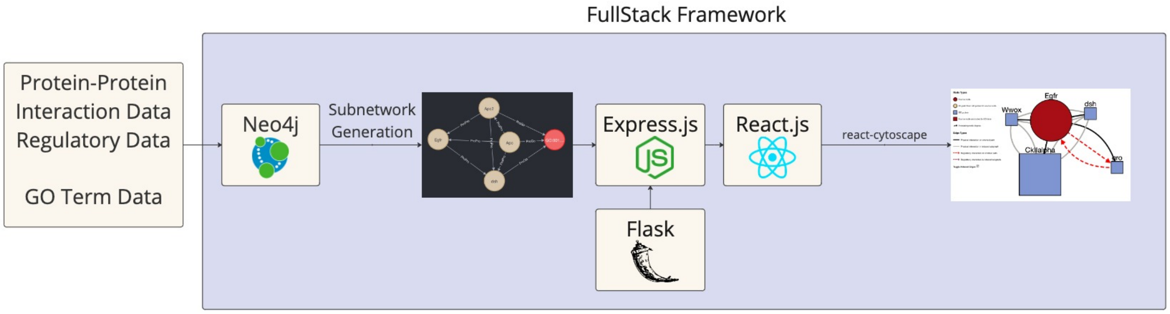

Figure S2: ProteinWeaver workflow and web architecture.

#### S3 Network Motif Enrichment

##### S3.1 Enrichment Scores for Mixed Motifs

Enrichment ratio scores, $\mathbb{E}$, are provided for each motif in the subnetwork, indicating whether the number of motifs observed in the subnetwork is surprising compared to the full interactome of a particular organism [29]. The score is calculated as the ratio of the motifs in the subnetwork to the expected number of motifs based on the structure of the entire network. Enrichment ratio scores greater than one suggest motif enrichment, while scores less than one indicate the motif occurs less frequently than expected. Given a graph $G = (V, E)$, a subgraph $G' = (V', E') \subseteq G$, and a motif $m$, let $C(G, m)$ be the count of motif $m$ in graph $G$. Let $|E|$ and $|E'|$ be the number of edges in the global network and subnetwork, respectively.

$$\mathbb{E} = \frac{C(G', m)}{\mu}, \text{ where} \tag{1}$$

$$\mu = \frac{|E'| C(G, m)}{|E|}. \tag{2}$$

Here, $\mu$ represents the expected number of motifs in the subnetwork, calculated by normalizing the number of edges in the subnetwork to the global network. This normalization accounts for uneven connectivity among nodes, as some nodes act as hubs with many connections while others have very few. This scaling ensures $\mu$ reflects the network's structure without bias and is derived based on a proportion using the calculation for subgraph concentration from Itzkovitz et al. [30]. A breakdown of the global counts of motifs $C(G, m)$ for each species is available in Figure S3.

To analyze the distribution of motifs, ProteinWeaver provides $Z$-scores, adapted from Kashtan et al.'s network motif detection method [31, 32], and an associated $p$-value for all enrichment scores. The $Z$-scores and $p$-value for a two-tailed test are calculated as:

$$Z = \frac{C(G', m) - \mu}{\sqrt{\mu}} \text{ and} \tag{3}$$

$$p = 2\left(1 - \Phi(|Z|)\right), \text{ where } \Phi(|Z|) = \frac{\left(1 + erf\left(\frac{|Z|}{\sqrt{2}}\right)\right)}{2}. \tag{4}$$

$\Phi(|Z|)$ is the cumulative distribution function (CDF) of the standard normal distribution, and $erf\left(\frac{|Z|}{\sqrt{2}}\right)$ is an error function based on the Abramowitz and Stegun [33] approximation error function. The error function provides a highly accurate approximation of the integral of the Gaussian function, giving context to the probability of observing a particular $Z$-score in a normal distribution.

#### S3.2 Motif Counts

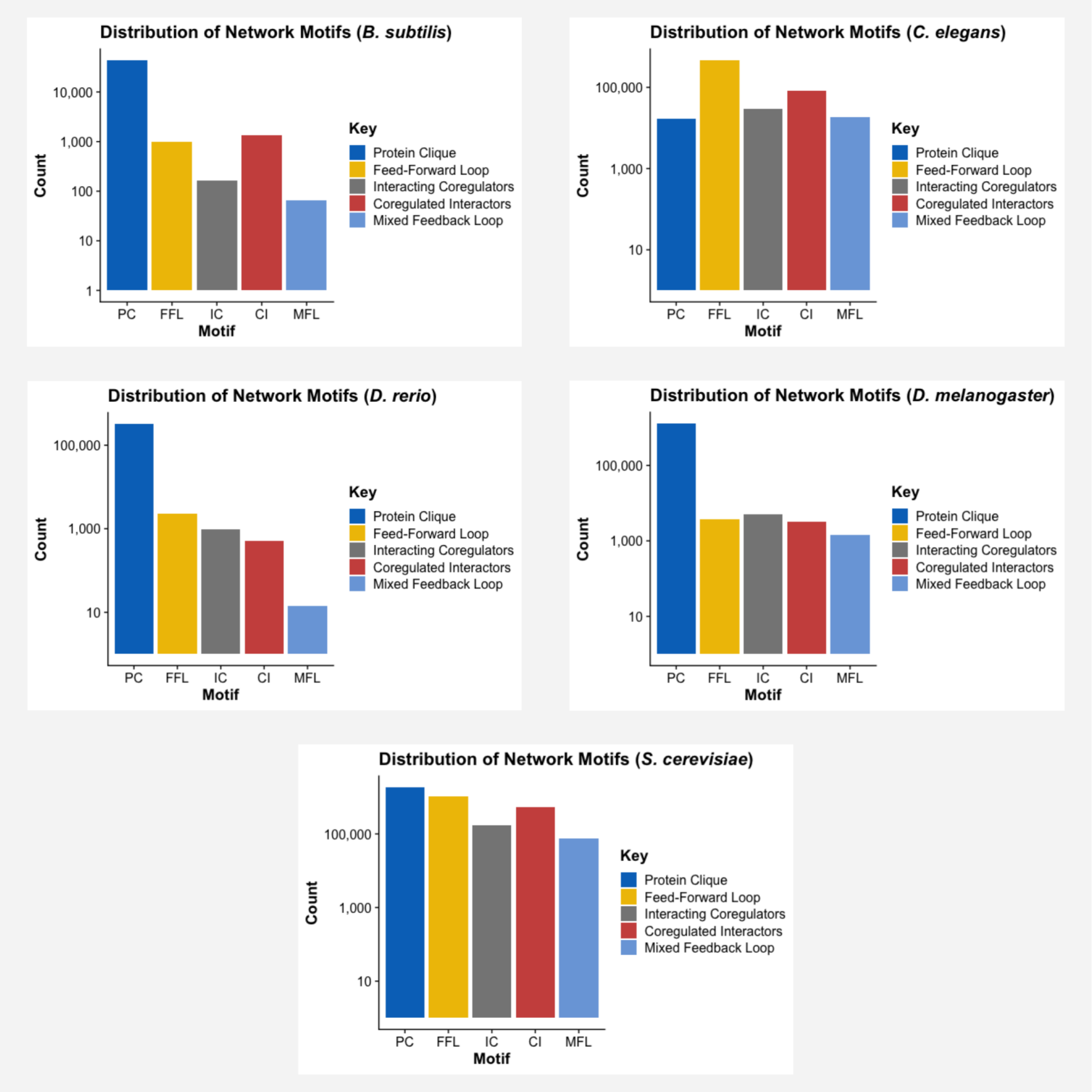

Figure S3: Global counts of each motif identified in ProteinWeaver by species.

### S4 GO Term Annotation Prediction

We ran personalized pagerank [34] as implemented by networkx on a bipartite graph of species-specific *ProGo* edges that include directly and indirectly-annotated proteins. Since indirect GO annotations are captured, the graph implicitly represents the GO hierarchy. PageRank restarts at one of the nodes annotated to the GO term (chosen uniformly at random from the set) with a damping factor $\alpha = 0.7$. The final score of the query node $s$ is the ranking of $s$ in the final visitation probabilities of the random walk, which corresponds to how near $s$ is to the proteins annotated to

GO term $t$. This can be pre-calculated for each species-specific *GoPro* graph.

##### S4.1 Comparator Methods

We developed various methods that could be categorized into four categories, where each category applied some similar method variant to assess potential GO term annotation prediction. All the methods are written in Python and can be accessed in the following repository:
https://github.com/Reed-CompBio/protein-function-prediction/tree/main/classes.

**Degree:** By simply looking at a given protein's outgoing edges (a combination of PPIs, regulatory interactions, and GO term annotations), calculate the score by the protein's degree.

**One-Hop GO Overlap:** This method assessed whether a protein should be annotated to a certain GO term by considering the neighbors of that protein and how many of those neighbors are annotated to the GO term. For a node $v$, let $N_v^{etype}$ be $v$'s neighbors constrained by the edge type *etype* (*ProPro*, *Reg*, *ProGo*, or *GoGo*). For a node $v$ and a GO term $t$ the One-Hop GO Overlap is calculated as:

$$score(v,t) = |N_v^{ProPro} \cap N_t^{ProGo}| \tag{5}$$

**Hypergeometric Distribution:** The hypergeometric $p$-value quantifies the probability that a node's neighbors are annotated to a GO term, accounting for the node degree and the number of GO annotations. We are given the graph $G = (V, E)$, a node $v$, and a GO term $t$. Like above, let $N_v^{etype}$ be $v$'s neighbors constrained by the edge type *etype* (*ProPro*, *Reg*, *ProGo*, or *GoGo*).

$$p(G,v,t) = \frac{\binom{K}{k} \cdot \binom{N-K}{n-k}}{\binom{N}{n}}, \text{ where} \tag{6}$$

$$N = \text{Total number of proteins in } G \tag{7}$$

$$K = v\text{'s protein neighbors: } |N_v^{ProPro}| \tag{8}$$

$$n = \text{One-Hop GO Overlap: } |N_v^{ProPro} \cap N_t^{ProGo}| \tag{9}$$

$$k = t\text{'s annotated proteins: } |N_t^{ProGo}| \tag{10}$$

##### S4.2 GO Term Annotation Prediction Evaluation

A benchmark pipeline was created to test how well various methods could score and predict a potential GO term annotation using PPI, regulatory interaction, and GO term annotations as inputs. The benchmark efforts, all data, methods, and figures can be reproduced in the following repository https://github.com/Reed-CompBio/protein-function-prediction. The following section outlines the steps of the pipeline.

The pipeline starts by first creating NetworkX representations of all the five major species used in ProteinWeaver using some configuration of PPI, regulatory interaction, and protein GO term annotations. These NetworkX variables are then used to easily access information about neighbors, a node degree, and the network scale that all the different algorithms will use to predict GO term annotations. Each of these algorithms is represented as a class in Python that has a standardized input and output. After the networks have been made, the workflow generates a data set of positive and negative samples, which will be used to calculate their ROC and PR curves later. Users can set a ratio of how many negatives should be generated for every positive. The algorithm to generate these datasets starts with the positive dataset, where we sample protein and go term edges in the network. Then, for an entry in the positive dataset, the algorithm uses a breadth-first search approach and finds the closest protein from the positive protein that also does not have an edge to the GO term whose degree is similar to the respective protein in the positive dataset; essentially creating a strong false pair of proteins and go term that has not been annotated yet but is close enough to the positive protein to be biologically relevant. Each GO term annotation prediction method will then use these datasets to calculate their respective scores on how well they can differentiate between the positive and negative datasets. We will use this data later to generate the figures. Finally, each method calculates a Receiver Operator curve and Precision/Recall curve to assess how each method performs. This pipeline is summarized in Figure S4.

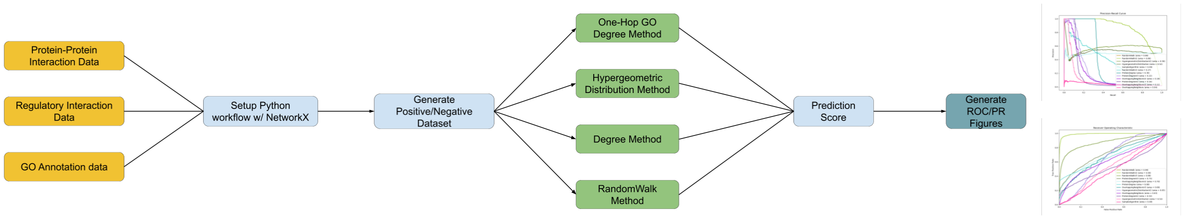

Figure S4: Pipeline for benchmarking the GO term annotation prediction. The pipeline starts with data collection; then, we set up a Python workflow by representing the data as NetworkX objects; next, we generate the list of positive and negative datasets; then, each GO term annotation prediction method runs all the data set in a standardized format; finally, we calculate the scores and generate the relevant figures.

#### S4.3 Complete Non-Inferred and Inferred Networks

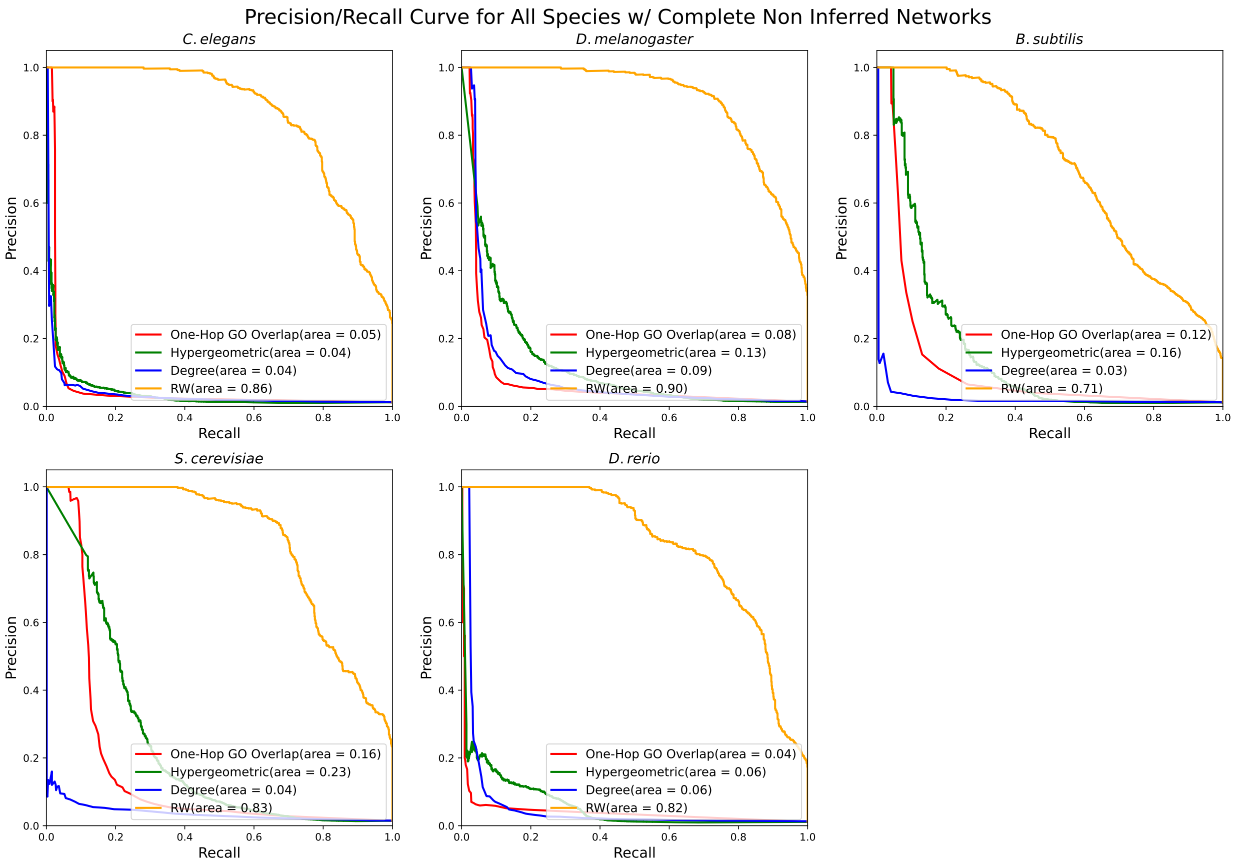

Figure S5: Precision/Recall curves for all species on the complete network without inferred annotations.

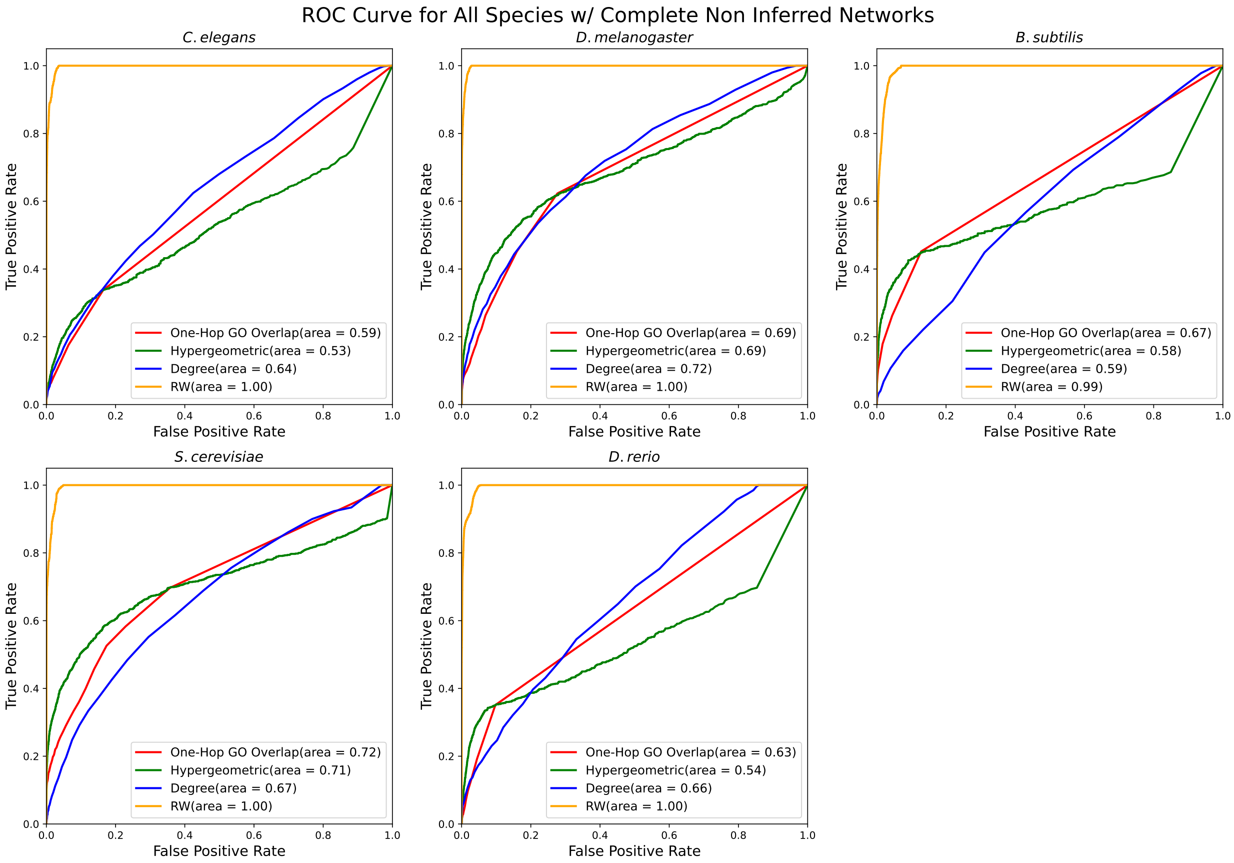

Figure S6: ROC curves for all species on the complete network without inferred annotations.

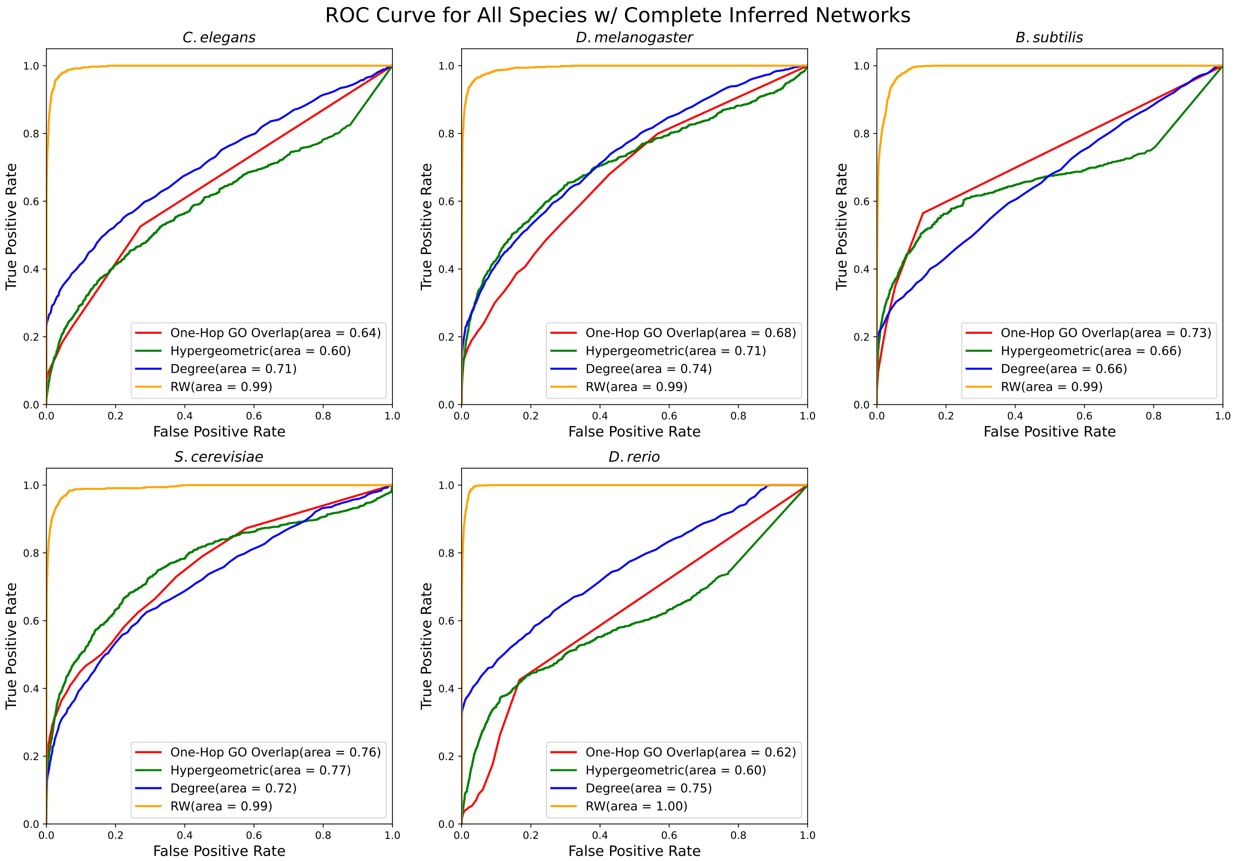

Figure S7: ROC curves for all species on the complete network with inferred annotations.

#### S4.4 RandomWalk on Various Network Parameters

We applied the RandomWalk method for all species on different subsets of the edges in the original graph: (1) the complete network with all edge types (the "Complete Inferred Network"), (2) the network with only direct *ProGo* edges (the "Non-Inferred Complete Network"), (3) a network consisting of only *ProGo* edges (the "Inferred ProGo Network"), and (4) a network consisting of only direct *ProGo* edges (the "Non-Inferred ProGo Network"). The random walk performs well on all scenarios for all species, with only subtle differences in the PR and ROC curves (Figures S8 and S9). Some species were consistently better or worse than others: for example *B. subtilis* had the lowest Precision/Recall value across all the configurations and *D. melanogaster* performed the best on non-inferred protein-GO networks.

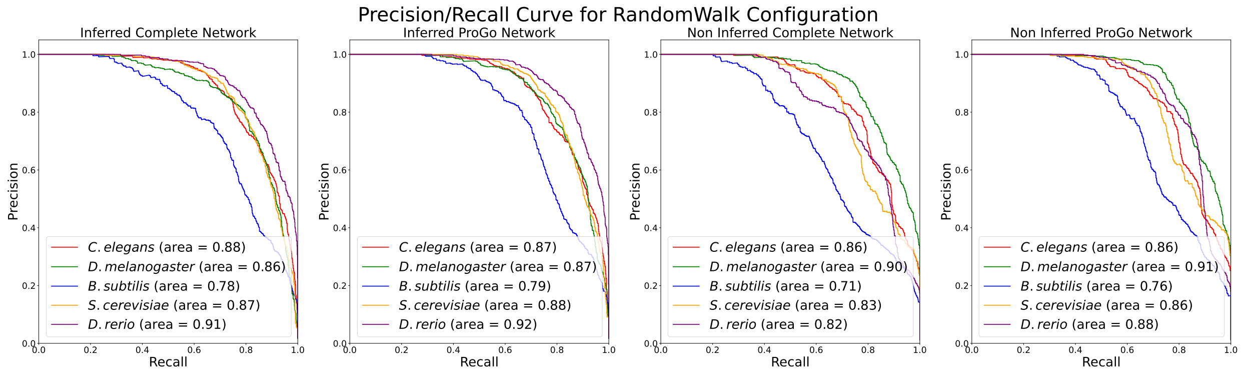

Figure S8: The RandomWalk prediction method in all five species across different background network configurations. Examples were sampled at a 1:100 positive-to-negative ratio.

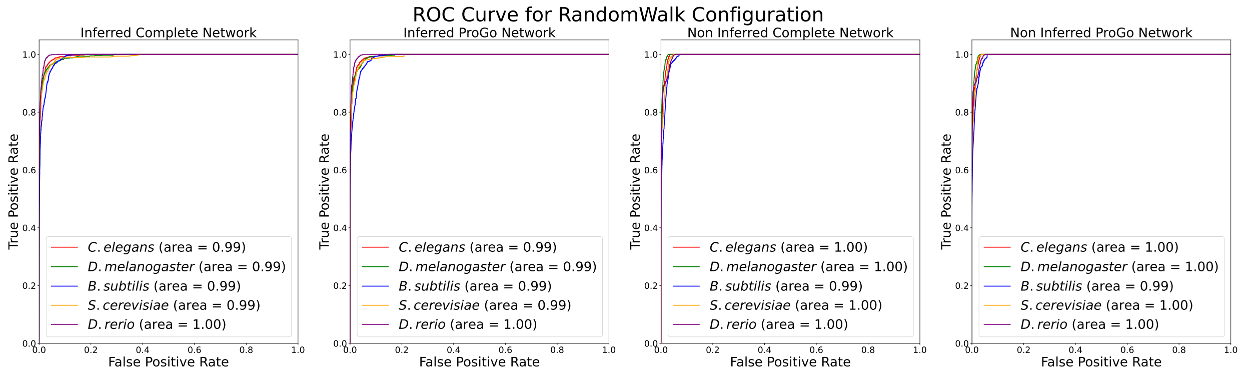

Figure S9: Four ROC curves that compare the RandomWalk method in all five species across varying network configurations. Examples were sampled at a 1:100 positive-to-negative ratio.

[5] Devin K. Schweppe et al. BioPlex Display: An Interactive Suite for Large-Scale AP–MS

Protein–Protein Interaction Data. *Journal of Proteome Research*, 17(1):722–726, January 2018.

[6] Christopher A Mancuso, Patrick S Bills, Douglas Krum, Jacob Newsted, Renming Liu, and Arjun Krishnan. Geneplexus: a web-server for gene discovery using network-based machine learning. *Nucleic Acids Research*, 50(W1):W358–W366, 2022.

[7] Thilakam Murali, Svetlana Pacifico, Jingkai Yu, Stephen Guest, George G Roberts, and Russell L Finley. Droid 2011: a comprehensive, integrated resource for protein, transcription factor, rna and gene interactions for drosophila. *Nucleic acids research*, 39(suppl_1):D736–D743, 2011.

[8] Tiago Pedreira, Christoph Elfmann, and Jörg Stülke. The current state of subti wiki, the database for the model organism bacillus subtilis. *Nucleic Acids Research*, 50(D1):D875–D882, 2022.

[9] Nicolas Alcaraz, Josch Pauling, Richa Batra, Eudes Barbosa, Alexander Junge, Anne GL Christensen, Vasco Azevedo, Henrik J Ditzel, and Jan Baumbach. Keypathwayminer 4.0: condition-specific pathway analysis by combining multiple omics studies and networks with cytoscape. *BMC systems biology*, 8(1):1–6, 2014.

[10] Jianguo Xia et al. NetworkAnalyst - integrative approaches for protein–protein interaction network analysis and visual exploration. *Nucleic Acids Research*, 42(W1):W167–W174, July 2014.

[11] David Binns et al. QuickGO: a web-based tool for Gene Ontology searching. *Bioinformatics*, 25(22):3045–3046, November 2009.

[12] Uniprot: the universal protein knowledgebase in 2023. *Nucleic Acids Research*, 51(D1):D523–D531, 2023.

[13] Seth Carbon et al. AmiGO: online access to ontology and annotation data. *Bioinformatics*, 25(2):288–289, January 2009.

[14] Paul W Sternberg, Kimberly Van Auken, Qinghua Wang, Adam Wright, Karen Yook, Magdalena Zarowiecki, Valerio Arnaboldi, Andrés Becerra, Stephanie Brown, Scott Cain, et al. Wormbase 2024: status and transitioning to alliance infrastructure. *Genetics*, 227(1):iyae050, 2024.

[15] Orsolya Liska, Balázs Bohár, András Hidas, Tamás Korcsmáros, Balázs Papp, Dávid Fazekas, and Eszter Ari. Tflink: an integrated gateway to access transcription factor–target gene interactions for multiple species. *Database*, 2022:baac083, 2022.

[16] Andy Zhao, Sophia Varady, Madelyn O'Kelley-Bangsberg, Vicki Deng, Amy Platenkamp, Petra Wijngaard, Miriam Bern, Wyatt Gormley, Elaine Kushkowski, Kat Thompson, et al. From network analysis to experimental validation: identification of regulators of non-muscle myosin ii contractility using the folded-gastrulation signaling pathway. *BMC Molecular and Cell Biology*, 24(1):32, 2023.

[17] Arzu Öztürk Çolak, Steven J Marygold, Giulia Antonazzo, Helen Attrill, Damien Goutte-Gattat, Victoria K Jenkins, Beverley B Matthews, Gillian Millburn, Gilberto dos Santos, Christopher J Tabone, and FlyBase Consortium. FlyBase: updates to the Drosophila genes and genomes database. *Genetics*, 227(1):iyad211, May 2024. ISSN 1943-2631. doi: 10.1093/genetics/iyad211.

[18] L Sian Gramates, Julie Agapite, Helen Attrill, Brian R Calvi, Madeline A Crosby, Gilberto Dos Santos, Joshua L Goodman, Damien Goutte-Gattat, Victoria K Jenkins, Thomas Kaufman, et al. Flybase: a guided tour of highlighted features. *Genetics*, 220(4):iyac035, 2022.

[19] Noemi del Toro et al. A new reference implementation of the PSICQUIC web service. *Nucleic Acids Research*, 41(W1):W601–W606, July 2013.

[20] ConvertCSV.com - Convert CSV To JSON, XML, SQL, ...

[33] Milton Abramowitz and Irene A. Stegun. *Handbook of Mathematical Functions*. Dover Publications, 1964.

[34] Lawrence Page. The pagerank citation ranking: Bringing order to the web. Technical report, Technical Report, 1999.
